# A Foraging-Theory-Based Model Captures the Spectrum of Human Behavioral Diversity in Sequential Decision Making

**DOI:** 10.1101/2025.05.06.652482

**Authors:** Dameon C Harrell, Meriam Zid, Veldon-James Laurie, Cathy S Chen, Nicola M Grissom, David P Darrow, R Becket Ebitz, Alexander B Herman

## Abstract

Decision-making tasks involving multiple, simultaneously presented options are mainstays of cognitive neuroscience and psychology and are increasingly important to the emerging field of computational psychiatry. Modeling approaches to these tasks overwhelmingly assume that participants make choices based on explicitly comparing the values of the presented options. Contrary to this long-held assumption, we found instead that humans employ a compare-to-threshold decision process, similar to theories of foraging, when making sequential decisions about concurrently available options. We confirmed this result in a large (1000 participant) dataset with multiple converging lines of evidence comparing both model fits and model generative performance. Value-comparison models were restricted to a reduced area of the potential space of single-trial outcome-dependent behavior, demonstrating an intrinsic limitation in the ability to reproduce strategy diversity. Furthermore, we found that using even the best-fit value-comparison model led to a substantial, systematic bias and a compression of individual differences in reconstructed behavior compared to the foraging-based model, leading to weaker predictions of behavioral health measures. Our results imply that studies using value-comparison models to link behavior with neural activity or psychiatric symptoms may be less sensitive to individual differences than a simple alternative based on ethological foraging.

## Introduction

Adaptive decision-making—balancing exploration of new opportunities with exploitation of known options—is fundamental to human behavior and frequently disrupted in psychiatric disorders [1]. Tasks probing these processes offer powerful tools for identifying individual differences in decision-making strategies. Exploration-exploitation impairments are consistently observed in conditions including depression, schizophrenia, autism, and chronic stress, making them valuable targets for computational psychiatry approaches that link computational-model parameters to psychiatric symptoms.

A critical challenge in this field is selecting tasks and models that accurately capture the full spectrum of human behavior—a prerequisite for understanding neurodivergent or pathological deviations. In this study, we examine the “restless bandit” task, which simulates dynamic reward environments to elicit realistic exploration-exploitation behavior. Most models of this task, for instance, reinforcement learning models, assume that people make their choices by tracking and comparing the values of all available options [2]. Although sophisticated formulations of these models have been developed to expand and improve their functioning, the foundational assumption of value-comparison-based choice is rarely questioned.

We recently developed an alternative “compare-to-threshold” model based on foraging theory [3], which assumes instead that individuals maintain a focal option and compare its value to an internal threshold to decide whether to exploit or explore. This approach offers computational efficiency (tracking only a single value and threshold regardless of option count) and conceptual alignment with psychiatric constructs through its threshold parameter, which reflects an individual’s perception of environmental reward potential. While similar models have been applied to patch-foraging tasks [4], they have not been applied to bandit tasks.

In a large sample (n=1000) performing a three-armed restless bandit task, we found that our foraging-based threshold model outperformed value-comparison approaches in explaining both the breadth and depth of human behavior. Furthermore, the foraging-based model’s parameters better captured psychiatric symptom variability than parameters from the best-fitting value-comparison models, providing additional evidence that a compare-to-threshold model aligns more closely with the underlying cognitive process.

## Results

### Task and Participant Behavior

We recruited 1000 gender-matched adults (500 males, 500 females) to participate in a restless 3-arm bandit task (Fig. 1A). Participants selected one of three playing cards by moving their mouse over their choice. Outcomes were binary: participants either received a point (win) or no point (loss). The probability of winning associated with each deck randomly drifted over time, encouraging participants to balance exploitation of good options with exploration for potentially better alternatives. Performance was evaluated by comparing the total number of rewarded trials to chance expectations. The vast majority of participants (925/1000) accumulated more rewards than would be expected by chance (see Methods for statistical model details).

**Fig 1.**
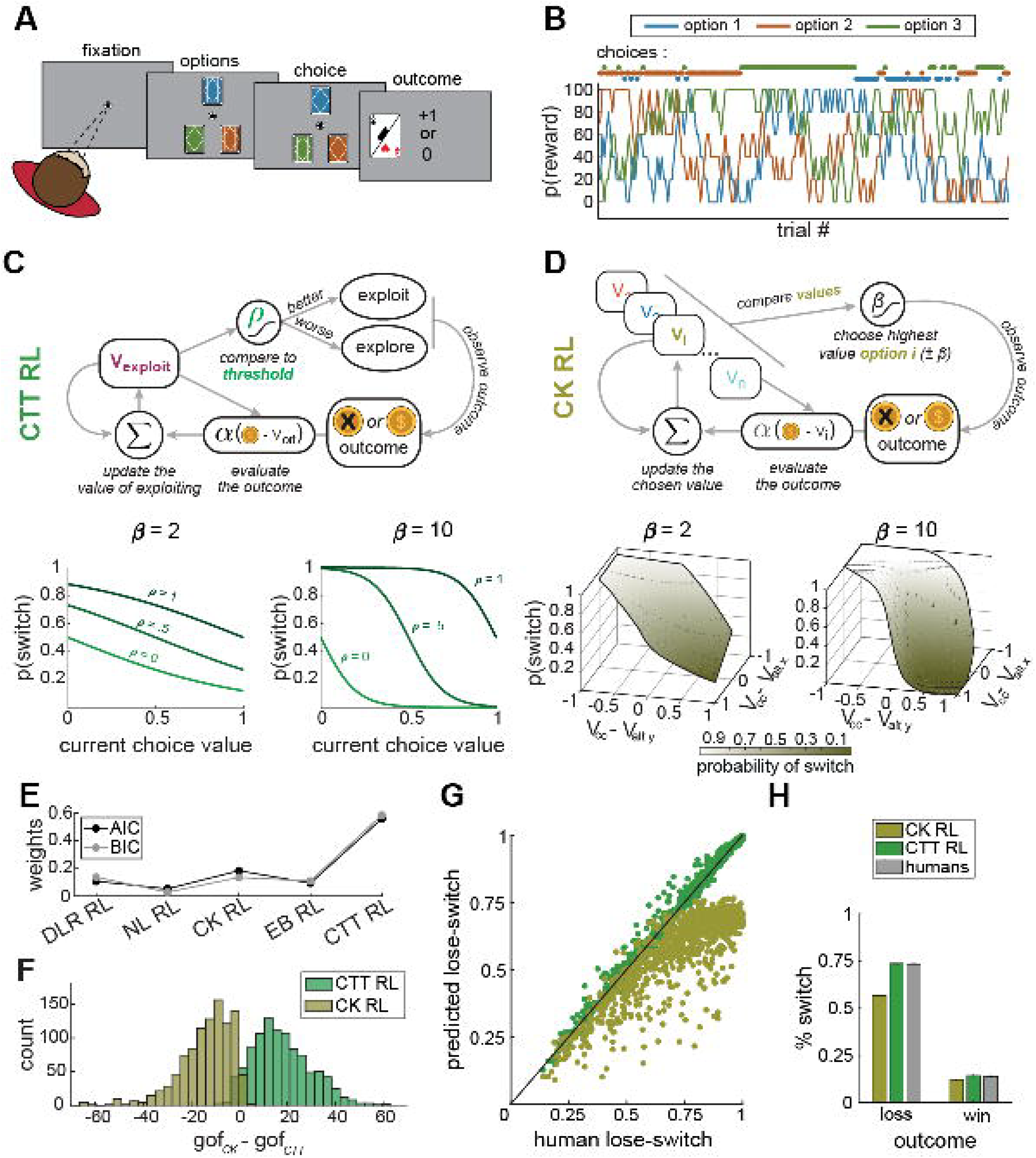
Task design, model comparison, and behavioral fits. (A) Three-deck bandit task interface. (B) Random-walk reward probabilities for each deck over time. (C) Foraging-RL model diagram (top) and switching probabilities at β = 2 and β = 10 (bottom), showing threshold-dependent behavior. (D) Simple RL model diagram (top) and switching probabilities as a function of value difference (bottom), illustrating greediness effects (β). (E) Model comparison using AIC/BIC weights shows foraging-RL outperforming all value-comparison models: ∼ 60% vs *<* 18% probability of being correct for the best value-comparison model). (F) Model recovery analysis showing limited overlap between foraging-RL and CK RL generated data fits, indicating distinct decision processes. (G) Actual vs. predicted lose-switch rates showing foraging-RL accurately predicts human behavior across the range, while CK RL fails above ∼ 0.7. (H) Average switch percentages following wins/losses for human participants and model simulations, demonstrating significant differences between human/foraging-RL data and CK RL predictions.

### Model Comparisons

To determine whether participants’ decision strategies more closely resembled a compare-to-threshold foraging-reinforcement learning (foraging-RL) model or one of several value-comparison RL models, we fitted each participant’s choice sequence using maximum likelihood estimation (MLE) [5]. We calculated Akaike’s Information Criterion (AIC) weights [6] and Bayesian Information Criterion (BIC) weights for model comparison. These weights reflect the probability that a particular model better explains the observed choices than all other candidate models [7], with BIC being more conservative for complex models by accounting for the number of observations.

The foraging-RL model provided the best fit for most participants, 578/1000 and 606/1000 according to AIC and BIC weights, respectively. In contrast, the best-fitting value-comparison model (Choice Kernel RL or CK RL) provided the best fit for only 181/1000 (AIC) and 136/1000 (BIC) participants (Table 1, Fig 1E). While the DLR (dual learning rate) RL model produced a slightly higher overall BIC weight (by 0.01) than the CK RL model, we selected the CK RL model to represent the value-comparison models for further analysis based on its larger AIC weight. This focus on one value-comparison model is justified since all value-comparison models share the same fundamental limitation compared to the foraging-RL model.

**Table 1.** Model Fits Comparison Average (across participants) raw AIC, AIC weights, raw BIC, and BIC weights. The table also contains the number of participants for whom the model was the best fit according to both AIC and BIC weights. Standard deviations are in parentheses.

| Model | Raw AIC | w(AIC) | n(AIC) | Raw BIC | w(BIC) | n(BIC) |
| --- | --- | --- | --- | --- | --- | --- |
| DLR RL | 204.7 (80) | 0.11<br>(0.23) | 113 | 215.8 (80) | 0.14<br>(0.27) | 150 |
| Noisy RL | 205.1 | 0.05 | 45 | 219.9 | 0.03 | 22 |
|  | (76.5) | (0.15) |  | (76.5) | (0.11) |  |
| CK RL | 199.6<br>(76.3) | 0.18<br>(0.33) | 181 | 214.5<br>(76.3) | 0.13<br>(0.3) | 136 |
| EB RL | 202.9 (83) | 0.09<br>(0.22) | 83 | 214 (83) | 0.11<br>(0.23) | 86 |
| foraging-RL | 185.3<br>(82.7) | 0.56<br>(0.45) | 578 | 196.4<br>(82.7) | 0.59<br>(0.45) | 606 |

### Model Recovery Analysis

The model comparison results above rely on predictive performance (how well a model predicts observed choices) and parsimony (favoring models with fewer parameters) [8].

However, these criteria only make relative comparisons—they identify which model is best among candidates without assessing absolute model quality. Additionally, models with equal parameter counts can differ in their flexibility to fit data. To address these concerns, we performed model recovery analysis to evaluate how well the models could mimic each other [8–10]. We used the Parametric Bootstrap Cross-Fitting Method (PBCM) to quantify each model’s ability to account for data generated by an alternative model [9]. When models cannot effectively mimic each other, this is evidence that they have distinct decision-making processes, and we can more confidently interpret differences between AIC and BIC weights. Figure 1F displays the distributions of differences in goodness-of-fit Δ*gof* = *gof*_*CK*_ − *gof*_*frl*_ between simulated data from the foraging-RL and CK RL models. Negative Δgof values indicate the CK RL model fit better, while positive values indicate the foraging-RL model fit better.

For data generated by the foraging-RL model (green bars), approximately 13.5% of simulations were better fit by the CK RL model. Conversely, for data generated by the CK RL model (gold bars), approximately 5% were better fit by the foraging-RL model. The limited overlap between these distributions suggests the decision processes of the CK and foraging-RL models are distinct, supporting the reliability of our AIC and BIC weight results, which favor the foraging-RL model.

### Generative Performance

While predictive performance provides valuable model comparison information, it is insufficient for falsifying particular models or measuring their capacity to produce behaviors of interest. Therefore, a model’s “generative performance”—its ability to reproduce key behavioral patterns—is also important to consider [8]. Figure 1G demonstrates the generative performance of both foraging-RL and CK-RL models in reproducing human switching behavior following losses in the bandit task. Each point (green for foraging-RL, gold for CK) represents the average lose-switch rate across 50 simulations (300 trials each) using parameters optimized for individual participants. The results clearly show that the foraging-RL model more accurately reproduces human switching behavior following losses compared to the CK RL model.

### Value-Comparison Model Dynamics and Limitations

To understand why value-comparison models struggle to reproduce switching behavior after losses, we analyzed the dynamics of the SoftMax decision rule across different value states with learning rate, α, set to 1. At this learning rate, an option’s value is immediately updated to match the most recent outcome (1 for win, 0 for loss), resulting in eight possible value states (2^3^) combinations for three options. Fig 2A presents state transition diagrams for value-comparison (w/ basic SoftMax) RL models with different inverse temperature (β) values. In these diagrams, *V*_*cc*_ represents the current choice’s value, followed by the values of the two alternative options. The left diagram (β = 8) shows “greedy” decision-making where the model strongly prefers higher-valued options. When the alternative options have zero value (state: *V*_*cc*_, 0, 0), the probability of switching after a win is zero. Switching after a win (win-switch) only occurs when at least one alternative option has a value of one. After a loss, the switching probability is 100%, when an at least one alternative option’s value is one (due to high β). But, once the model reaches state (*V*_*cc*_, 0, 0), it cannot return to states where alternatives have non-zero values because it will not switch after a win. This limits the maximum lose-switch probability to 67%. The right diagram (β = 2) shows less deterministic decision-making. Here, the win-switch probability in state (*V*_*cc*_, 0, 0) increases to 21%, allowing the model to occasionally reach states where alternative options have non-zero values after losses.

**Fig 2.**
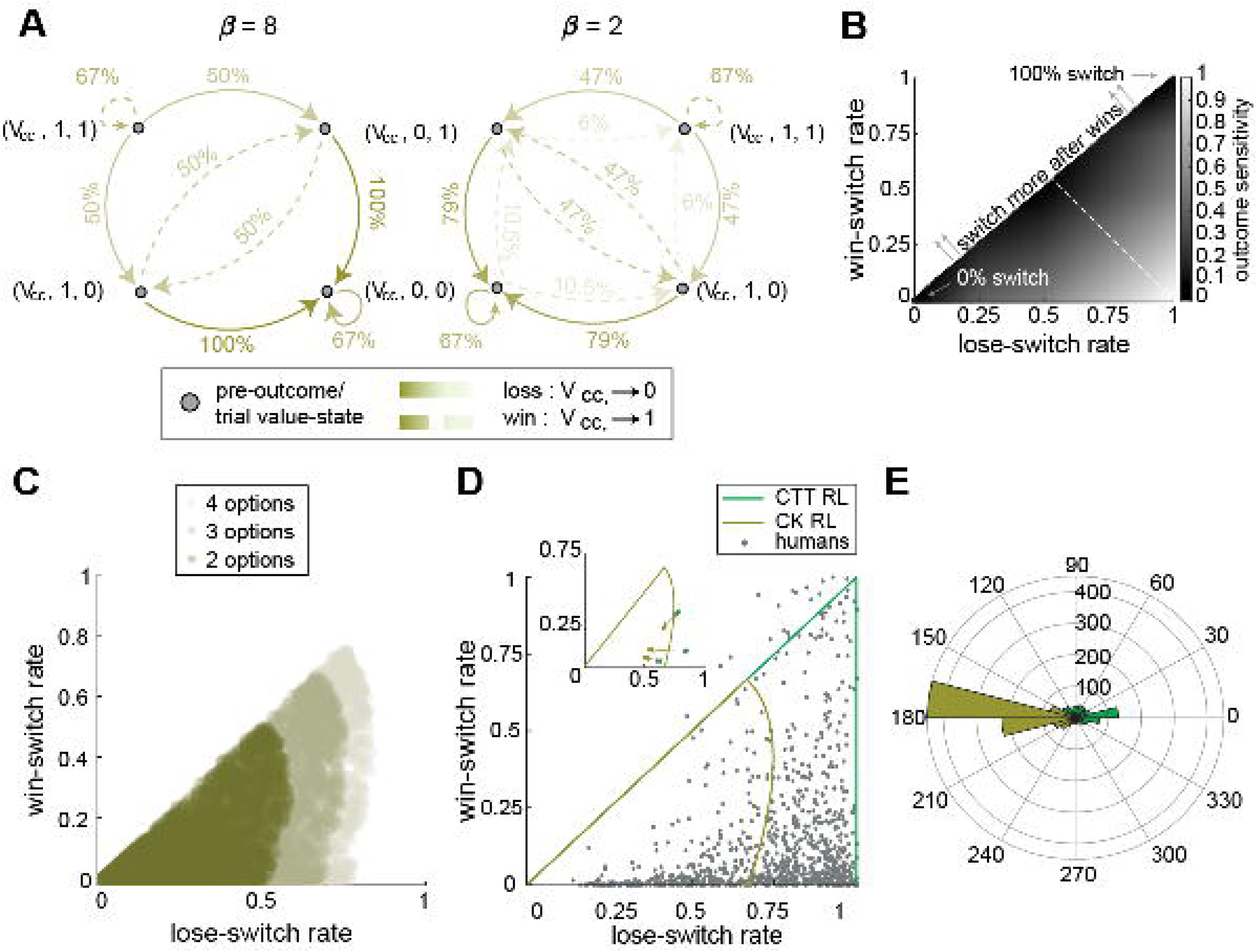
Value-Comparison Model Limitations in Outcome Sensitivity. (A) Value-state transition diagrams for SoftMax decision rule (α = 1) at two different β values. With β = 8 (left), “greedy” decisions eventually trap the model in state “*V*_*cc*_,0,0” limiting lose-switch probability to 67%. At β = 2 (right), increased stochasticity allows occasional win-switches, enabling higher lose-switch rates. (B) Outcome sensitivity space with win-switch rates (vertical) and lose-switch rates (horizontal). The shaded region below the diagonal represents lose-switch % *>*= win-switch % behavior; corners represent extreme sensitivities/insensitivities. (C) 10,000 trial simulations of CK RL for 2, 3 and 4 armed bandit task demonstrating limitations in outcome sensitivity//lose-switch rate dependency on number of values to compare. (D) Human participants’ outcome sensitivity (gray dots) with behavioral ranges producible by foraging-RL (green outline) and CK RL (gold outline) models. Inset shows how CK RL best-fit simulation bias outcome sensitivities towards lower lose-switch rates even when the original participant’s outcome sensitivity is within CK RL’s reproducible region. (E) Polar histograms of difference vectors between participant and model-simulated sensitivities, showing foraging-RL’s evenly distributed errors versus CK RL’s systematic bias toward underestimating lose-switch rates.

In summary, the SoftMax decision rule creates an inherent trade-off: higher lose-switch rates require higher win-switch rates, which depend on β. When β is too high, lose-switch rate is capped at 67%. When β = 0 (completely random), the lose-switch rate is also 67%. The maximum lose-switch rate occurs at an intermediate β value but remains below 100% due to the required stochasticity. Additionally, the limitations of lose-switch and win-switch rates are determined by the number of values compared within the SoftMax (Fig 2C).

### Outcome Sensitivity Framework

We conceptualize these limitations using “outcome sensitivity”—how responsive an agent is to wins or losses. We define a two-dimensional outcome sensitivity space with loss-switch rates on one axis and win-switch rates on the other (Fig. 2B). This space has several important reference points: maximal outcome sensitivity (lose-switch = 100%, win-switch = 0%, also known as a win-stay/lose-switch strategy) appears in the lower right corner; maximally biased insensitivity (either never switching or always switching regardless of outcome) appears in the lower left and upper right corners; and unbiased insensitivity (random 50% switching probability) appears at the midpoint (0.5, 0.5). The area below the diagonal (lose-switch *>* win-switch) is most relevant for reward-maximizing behavior. The lower right corner has particular relevance as an anchor point because a pure win-stay/lose-switch (also known as Pavlov) strategy is a simple heuristic solution to a wide range of decision-making problems ([11]; [12]; [13]), including bandits ([14]). An individual point in this space thus represents the degree to which that participant’s behavior deviates from win-stay/lose switch along dimensions of sensitivity to loss and sensitivity to wins.

### Human and Model Behavior in Outcome Sensitivity Space

Figure 2D displays human participants’ behavior in outcome sensitivity space (gray dots).

The vast majority (98.3%) showed higher lose-switch than win-switch rates, with 64.8% having win-switch rates below 10%. Over a quarter (252/1000) had win-switch rates below 10% and lose-switch rates above 80%. The gold and green outlines in Fig. 2D show the behavioral ranges that the CK RL and foraging-RL models can produce. The foraging-RL model covers the entire region where lose-switch rates exceed win-switch rates. In contrast, the CK RL model covers only a portion of this space. Its maximum win-switch rate is 67% (with completely random selection, β = 0). When win-switch is 0%, the lose-switch rate is limited to 67%. Higher lose-switch rates are only possible with increased win-switch rates. Figure 2E uses polar histograms to visualize differences between human and model-simulated behavior. Specifically, it displays the distributions of the angles of the vectors formed by the line segments connecting each human participant’s point in outcome sensitivity space to the point of the corresponding best-fit optimized foraging-RL and CK RL simulations. The angles of foraging-RL difference vectors show a relatively even distribution around the polar plane, indicating no systematic bias. In contrast, the CK RL difference vector angles cluster between 165-195 degrees, with 673 out of 1000 simulations underestimating participants’ lose-switch rates more than they misestimate win-switch rates. The magnitude of these vectors further confirms foraging-RL’s superior generative performance. Most foraging-RL simulations (695/1000) are within 0.02 switches/outcome of human data, with none exceeding 0.18. CK RL simulations show a more uniform distribution of differences, with some as large as 0.65 switches/outcome, and nearly half (484/1000) exceeding the maximum foraging-RL difference. The magnitude is calculated as the Euclidean distance between the true and simulated outcome sensitivity points (Eq. (1)).

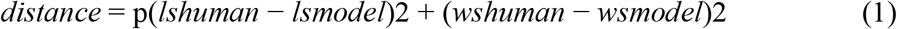

### Outcome Sensitivity in Model Parameter Space

We defined a unified outcome sensitivity measure as: (lose-switch rate - 0.5) + (win-stay rate - 0.5), where win-stay = 1 - win-switch. This yields: maximum sensitivity at lose-switch = 1, win-stay = 1 (outcome sensitivity = 1); unbiased insensitivity at lose-switch = 0.5, win-stay = 0.5 (outcome sensitivity = 0); and maximum biased insensitivity when one rate is 1 and the other is 0 (also outcome sensitivity = 0). We then examined this measure across the different models we considered.

### Simple RL Model

For the simplest value-comparison RL model (basic delta learning + basic SoftMax) (Fig. 3A), when β *>* 2, outcome sensitivity primarily depends on α, showing a clear gradient from high to low sensitivity as α decreases. When β *<* 2, outcome sensitivity is mainly a function of β and is relatively independent of α. Outcome sensitivity reaches a maximum of approximately 0.68 and is minimized when either β or α equals zero.

**Fig 3.**
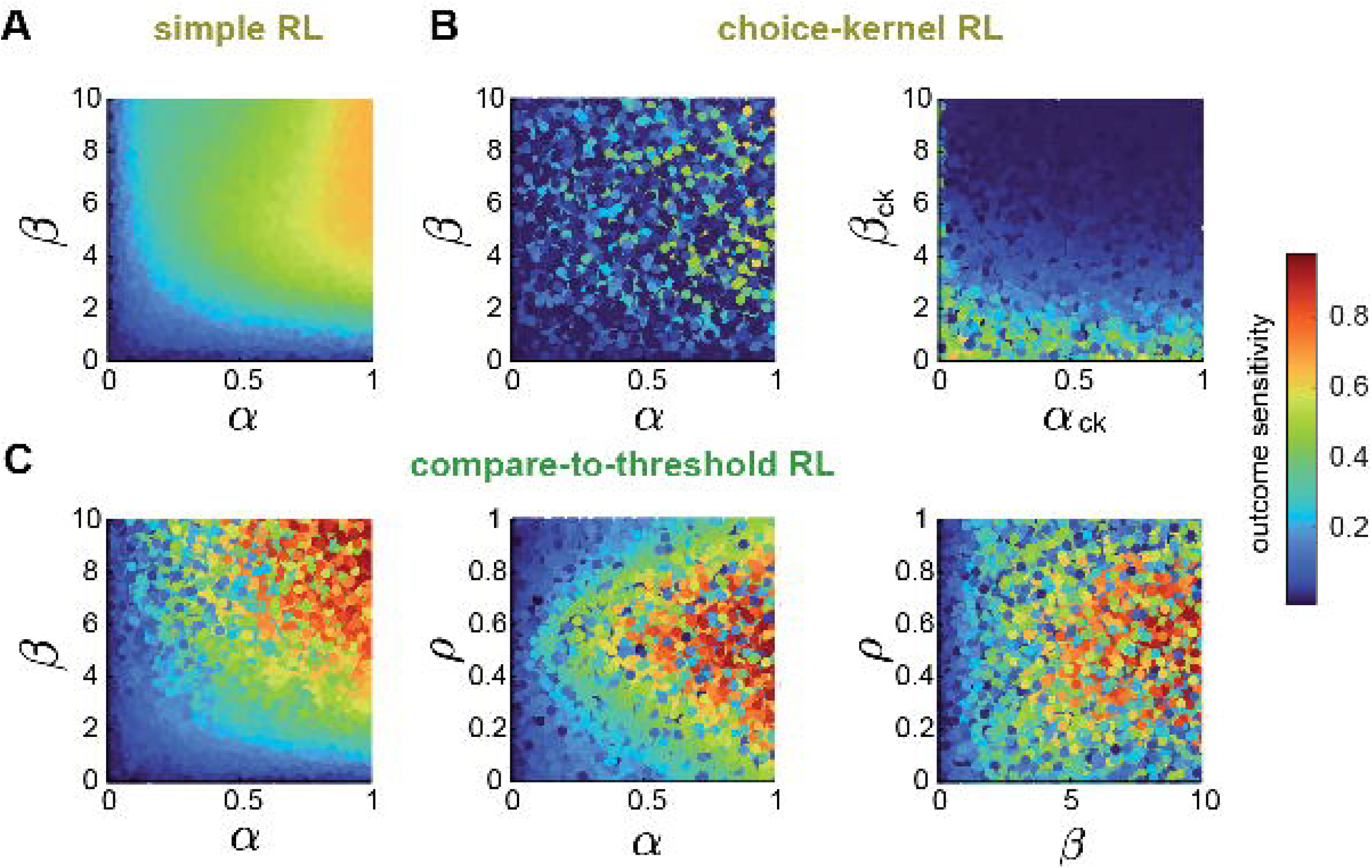
Outcome Sensitivity in Model Parameter Space. (A) Outcome sensitivity as a function of α and β parameters, for the most basic value comparison RL model, is primarily driven by α when β *>* 2. (B) Outcome sensitivity for the CK RL model shows less dependency on α and less overall sensitivity (left). CK RL’s overall reduction in outcome sensitivity, compared to Simple RL, is mainly due to its β_*ck*_ parameter, which leads to biased insensitivity as it increases. (right) (C) foraging-RL’s outcome sensitivity appears similar to Simple RL in α,β space (left), however, a more heterogeneous mixture of sensitivities is possible when α *>* .5. Foraging-RL’s ρ parameter allows for biased insensitivity even with higher values of α (center). Foraging-RL outcome sensitivities are roughly symmetric around ρ = .5 (center and right panels) regardless of outcome. Outcome sensitivity appears inversely proportional to β_*ck*_ when β_*ck*_ *<* 3 but shows little dependency on α_*ck*_.

### CK RL Model

The CK RL model (Fig. 3B left) shows a more complex parameter landscape with no clear gradient in outcome sensitivity relative to α and β. Low outcome sensitivities (*<* 0.2) appear throughout the parameter space, even with high α and β values. The highest sensitivities occur with high α and β, but the choice kernel parameters (α_*ck*_ and β_*ck*_) often drive behavior toward biased insensitivity. Figure 3B (right) shows that high β_*ck*_ values (*>* 5) generate consistently low outcome sensitivities (*<* 0.2) by biasing behavior toward repeatedly choosing the same option.

### Foraging-RL Model

The foraging-RL model’s outcome sensitivity (Fig. 3C left) resembles the Simple RL model with important differences. When β *>* 2, outcome sensitivity primarily depends on α, but unlike Simple RL, the relationship is not monotonic as α decreases. Outcome sensitivity varies from 1 to 0.4 even with high β(*>* 6) and high α, ∼ (0.8 − 1). Foraging-RL can produce biased outcome insensitivity with α as high as 0.5, while Simple RL requires α *<* 0.2. Figure 3C (center) shows outcome sensitivity as a function of α and threshold parameter ρ. Outcome sensitivity is roughly symmetric about ρ = 0.5, decreasing as ρ moves away from 0.5 in either direction. The range of possible sensitivities narrows as α decreases, forming a triangular pattern in α-ρ space. The threshold ρ functions as a bias term that decouples sensitivity to wins from sensitivity to losses. This allows the model to achieve maximum sensitivity to both simultaneously, which is impossible for value-comparison models using the SoftMax decision process, where sensitivity to losses is inherently tied to sensitivity to wins.

### Prediction of behavioral measures

We hypothesized that foraging-RL’s superior ability to capture individual differences in task behavior would extend to better predictions of personality and mental health symptom measures. We analyzed a range of such self-report measures, covering depression, anxiety, pain, sleep quality, anhedonia, motivation, autism-like traits, and cognitive flexibility. Models predicting these measures from the foraging-RL parameters outperformed CK RL based models on 10/13 of these scales by AIC weight (Fig. 4B). Next, we examined two of these domains in depth, apathy (behavioral motivation) and anxiety, which we have recently shown to co-vary with explore-exploit behavior on this task [15]. The foraging-RL model better accounted for variance in both symptom domains. The foraging-RL parameters also provided a more parsimonious description of the cognitive processes that are related to apathy and anxiety. For both symptoms, the explained variance was concentrated in one foraging-RL parameter, the foraging threshold ρ, while explained variance was more evenly distributed across the parameters of the CK RL model. Furthermore, the foraging-RL ρ parameter explained more variance in both symptom measures than all the CK RL parameters combined (Fig. 4A).

**Fig 4.**
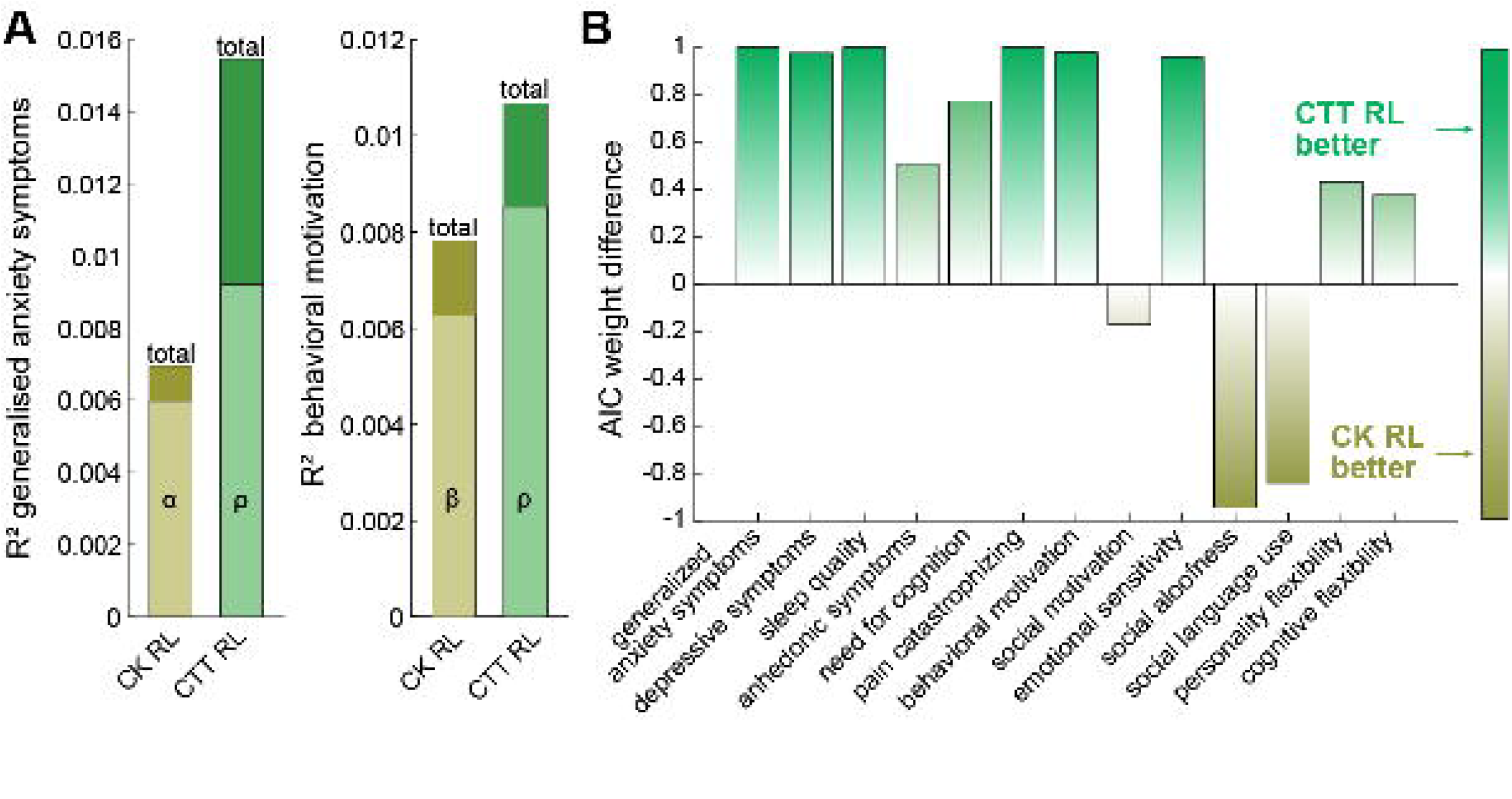
Comparing the predictive utility of CK RL and foraging-RL models for anxiety, depression, and other self-reported measures. (A) Proportion of variance (*R*^2^) in generalized anxiety (left) and depressive (right) symptoms explained by CK RL and foraging-RL models. For each model, the total variance explained is further broken down into the top contributing parameter from each model (α for CK RL; and ρ for foraging-RL). (B) Akaike weights for full parameter models, reflecting the relative likelihood of either the foraging-RL or CK RL model, for a range of self-reported measures. Bars above zero indicate the probability, given the data and parameters and set of models, that the foraging-RL based model is the correct model; bars below zero indicate the CK RL model has the higher probability of being the correct model.

## Discussion

Our findings provide evidence that humans utilize a compare-to-threshold, foraging-like decision process rather than a value-comparison approach when making sequential decisions about concurrently available options. This represents a fundamental shift in our understanding of human decision making, with significant implications for computational psychiatry and cognitive neuroscience.

The superior performance of the foraging-RL model over traditional value-comparison RL models in explaining human behavior stems from its ability to capture a much broader range of decision strategies. The limitations of value-comparison models are not merely implementation details but reflect fundamental constraints in their architecture. Specifically, the SoftMax decision rule used in these models creates an intrinsic trade-off between win-switch and lose-switch rates, leading to model value-states with bounded lose-switch or win-switch rates. These bounds prevent value comparison models from populating the full outcome-sensitivity lose-switch *>* win-switch subspace, restricting them to a portion of this subspace instead. As a result, when fit to real human behavior, value-comparison models compress the observed behavior into this limited region, obscuring individual differences and introducing systematic biases. This compression implies that the parameters from even the best-fit value-comparison RL model must be interpreted with caution. In contrast, the foraging-RL model’s threshold parameter allows for independent modulation of win-switch and lose-switch responses, enabling it to capture the full range of possible strategies and thus reproduce the full range of observed human behavior.

Our findings have important implications for computational psychiatry, underscoring the importance of accurate modeling of individual differences [16]. The constrained outcome-sensitivity and systematic bias towards lower lose-switch rates introduced by value-comparison models compresses individual differences and may obscure or even misconstrue important relationships between decision-making parameters and psychiatric symptoms. The inability of these models to represent agents with high lose-switch rates and low win-switch rates—a pattern we observed in many human participants—could lead researchers to mischaracterize psychiatric populations or miss important correlations between model parameters and clinical symptoms. In contrast, the coverage of outcome-sensitivity space produced by the foraging-RL model resulted in a superior ability to capture individual differences in a diverse range of symptom measures. In two important symptom domains, apathy (behavioral motivation) and anxiety, a single parameter from the foraging-RL model (the threshold) explained more variance than all of the parameters combined from the best-fitting value-comparison RL model. This parsimony offers significant advantages for inferring the connections between symptoms and cognitive processes and, ultimately, putative physiological mechanisms [17].

The threshold parameter in the foraging-RL model may have special relevance for psychiatric research as it may relate to an individual’s perception of environmental reward, as it does in classic foraging theory ([3]; [18]). Altered threshold values could represent different expectations about reward availability—a construct directly relevant to various neuropsychiatric disorders where reward expectation is affected. For instance, we found that thresholds decrease as apathy increases, implying that an individual’s overall level of motivation is related to how rewarding they see the environment. In contrast, thresholds increased with anxiety, potentially reflecting a response to risk aversion or the worry that unselected options may be better than the choice that was made.

Future research should explore whether threshold differences can differentiate clinical populations and whether these parameters are more sensitive to treatment effects than those derived from value-comparison models.

Several limitations of the present study should be addressed in future work. First, while our task uses a “restless bandit” paradigm to encourage exploration-exploitation decisions, other decision-making contexts might elicit different strategies. Second, our foraging-RL model utilized a constant threshold, but implementing a dynamic threshold that updates based on reward history could further improve model fit and better represent adaptation to changing environments. Third, although we examined a wide range of reinforcement learning models, there may be other possible formulations that would overcome the limitations that seem to be inherent to value comparison.

It is worth noting that value-comparison models have been foundational in computational neuroscience, leading to important insights about dopaminergic prediction error signals and reward learning ([19]; [20]). Because prediction error-based learning is central to both value-comparison and compare-to-threshold models, our results suggest that some of these past findings may warrant re-examination from the perspective of foraging theory. In our foraging-based model, the prediction refers to the value of the decision, stay or switch, rather than the values of the choices. This distinction may yield new insights into how value signals and thresholds are represented in the brain.

Our results challenge a fundamental assumption in decision-making research and offer a more ecologically valid framework for understanding how humans navigate exploration-exploitation dilemmas. The compare-to-threshold perspective provides superior explanatory power for individual differences in decision strategies and may ultimately advance our understanding of decision-making impairments in psychiatric disorders.

## Methods

This is a new analysis of a previously published dataset (Yan et al. 2025).

### Participants

The required number of participants was determined through a priori power analysis. To detect correlations of r = 0.1 with symptom measures (typical for individual differences research) with 80% power at α = 0.05, we required a minimum sample of 782 participants. We recruited 1500 participants to account for expected exclusions based on previous large-sample online studies ([21]; [22]) and our own pilot work, expecting to achieve a sample of between 900-1100 participants, thus allowing for a buffer above the minimum sample size. Our final sample of 1000 participants (500 female) provided 98% power to detect r = 0.1. We recruited a sample of 1512 participants (non-clinical sample) via Prolific (Prolific. co); exclusion criteria included current or history of neurological and psychiatric disorders. Participants were excluded if they did not complete all questionnaires (3.57% of initial sample) or they did not complete the bandit task (30.22% of initial sample). 1000 participants completed all questionnaires and the restless bandit task (age range 18-54, mean ± SD = 28.446 ± 10.354 years; gender, 493 female). All participants were compensated for their time in accordance with Prolific’s minimum wage.

### Task

Each participant completed a 300-trial, 3-arm restless bandit task. On each trial, participants were presented with images of 3 face-down playing cards, each associated with a deck of a specific color (arm) and instructed to choose one by moving the cursor over their choice. Following every choice, participants received feedback: either 1 point (a win) or no points (a loss). The probability of winning on any trial was independent for each deck and the product of a “random walk”. There was a 67% probability that on any trial a given deck might increase or decrease its win yield rate by an increment of 10%. Initial win yield rates for each deck were randomized, and win rates could vary from 100% to 10%.

### Raw Performance Assessment

The number of wins expected by chance for each participant’s reward schedule (win yield rates across all decks over 300 trials) was determined by generating distributions (probability mass functions; PMF) of the number of winning trials accumulated by randomly selecting a deck on each of the 300 trials for that participant’s reward schedule. Specifically, 1000 300-trial simulations were used to produce each participant’s PMF for chance expected number of wins. The number of winning trials actually accumulated by each participant was then used to determine the probability of accumulating that number of wins by chance according to the PMF. If the probability was less than .05, the participant’s number of wins was assessed as being non-chance.

### Model Details

#### Compare-to-Threshold foraging-RL

Foraging-RL is a model comprising a learning rule Eq (2), for estimating option values, and a decision rule. In contrast to the more common version of the delta learning rule (see Eq (5), only the value of the currently selected option is maintained and the value resets to .5 after a decision to switch to an alternative option. The learning rate α (∈ [0,1]) determines the rate at which the current outcome, 0(*t*), is incorporated into the value estimate, *v*(*t*):

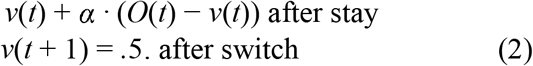

The foraging-RL model uses a compare-to-threshold decision rule, Eq (3). The single maintained value is compared, via a difference in the exponential, to a threshold, ρ (∈ [0,1]), a parameter of the model. β (∈ **R**^+^), another model parameter, adjusts sensitivity to the difference between *v*(*t*) and ρ, thus partially determining stochasticity in the decision process. Our foraging-RL decision rule outputs the probability of switching, not the probability of choosing a *particular* option. When the model decides to switch, it chooses between the previously unselected alternatives with equal probability.

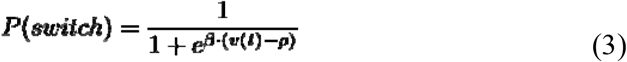

#### SoftMax (value-comparison) Decision Rule

The primary difference between the foraging-RL and the value-comparison RL models is found in their decision rules. All of the value-comparison models in our study use the SoftMax equation Eq (4), aka the Boltzmann distribution [23], either in its basic (DLR RL) or a modified form (CK, EB, and Noisy RL). The output of the SoftMax is the probability of choosing a particular option. More precisely, the SoftMax model’s decisions of “non-greedy” choices are based on the relative values of the available options. Here, greedy is defined as always selecting the option with the highest value. The right-hand side of the top portion of Eq. (4) can be rearranged to see the value comparisons (differences) more explicitly. This rearrangement is shown in the bottom half of Eq. (4). *P*[*C*(*t,i*) = 1] is the probability of choosing option i on trial t; *C*(*t,i*) = 1 if option i was selected on trial t, *C*(*t,i*) = 0 otherwise. β (∈ **R**^+^) (aka “inverse temperature”) is commonly thought of as controlling the am noise in the decision process and therefore determines its stochasticity; β also determines the sensitivity to value differences. When β is zero, the decision process is insensitive to value difference, completely random, and chooses uniformly between each option. As β approaches positive infinity, the SoftMax output becomes “greedy”: it always selects the option with the highest value.

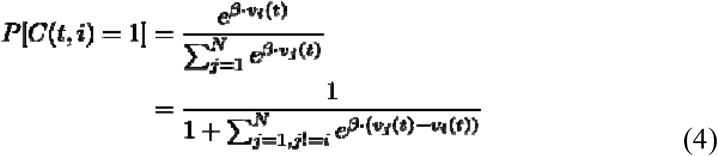

#### Choice Kernel (CK) RL

The CK RL model [5] employs two learning rules and uses a modified SoftMax for its decision rule. The basic delta learning rule, Eq (5) [23,24], updates the value of options. Learning rate α (∈ [0,1]) determines the rate at which the current outcome, 0(*t*), is incorporated into the value estimate, *v*_*j*_(*t*). *O*(*t*) = 1 for a win or *O*(*t*) = 0 for a loss. Only the value of the chosen option is updated during a trial; δ_*j*_(*t*) = 1 if option j was chosen during trial t and is 0 otherwise. An additional learning rule is used to maintain an estimate of how frequently each option is chosen in order to account for the tendency of individuals to repeat their previous actions [5]. Choice frequency estimates are updated on each trial according to Eq (6). *ck*_*j*_(*t*) is the estimated choice frequency, α_*ck*_ (∈ (0,1]), is the choice kernel learning rate. *C*(*t,j*) = 1 if option j was selected at trial t, otherwise *C*(*t,j*) = 0. Unlike expected value, choice kernel values for *all* options are updated on every trial.

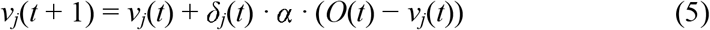

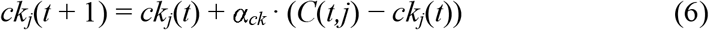

The choice kernel modified SoftMax, Eq (7), extends the basic SoftMax of Eq (4) with an additional term in the exponents consisting of the product of a second inverse temperature, β_*ck*_ (∈ **R**^+^), and choice frequency estimates from Eq (6). Larger β_*ck*_ values in combination with higher choice frequencies will increase the probability that an option is chosen repeatedly. If β_*ck*_ is set to zero this rule reverts to the basic SoftMax Eq (4).

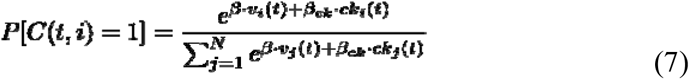

#### Dual Learning Rate (DLR) RL

DLR RL was inspired by the “optimism bias”, which suggests humans respond asymmetrically to rewards and punishment [25]. This model’s learning rule comprises a set of equations, Eq (8). This learning rule is similar to the delta learning rule (Eq (5)) except the learning rate, α (∈ [0,1]), depends on the outcome of the current trial.

If the outcome is a loss (*O*(*t*) = 0), a learning rate associated with losses is used to update the value. If the outcome is a win (*O*(*t*) = 1), a learning rate associated with wins is used to update the value. The DLR RL model uses the basic SoftMax, Eq (4), to model its decision process.

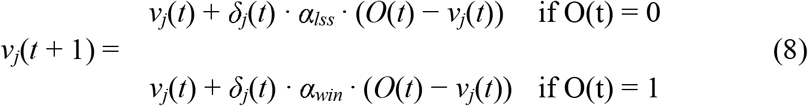

#### Noisy Learning RL

The Noisy Learning RL model, derived by Findling and Colleagues [26], employs a learning rule, Eq (9), similar to the delta learning rule in Eq (5). However, its learning rule adds a time-varying noise term, ε(*t*). ε(*t*) is a zero-mean, normally distributed noise component, ε(*t*) ∼ N(0,σ(*t*)). The standard deviation, σ(*t*), is equal to a constant fraction of the prediction error magnitude Eq (10). ζ (∈ [0,.75]) is a free parameter of the model. When ζ = 0, this rule is equal to the delta learning rule of Eq (5).

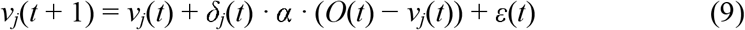

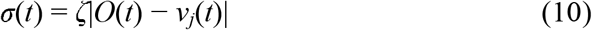

The Noisy RL model uses the decision rule presented in Eq 11. It is the basic SoftMax in Eq (11) with an added “choice hysteresis” term in the exponential. The choice hysteresis ξ (∈ [0,1]) is a free parameter of the model and serves a similar purpose to the choice kernel in Eq (7) ^1^; it increases the probability of an option being chosen during the current trial if it was chosen on the previous trial. Specifically, δ_*j*_(*t* − 1) = 1 if option j was chosen on trial t-1 and is 0 otherwise and δ_*i*_(*t* − 1) = 1 if option i was chosen on trial t-1 and is 0 otherwise.

If ξ is set to zero this decision rule reverts to the basic SoftMax. Note the equation as presented here is slightly modified from how it is written in Findling et. al [26] by using a delta function for the choice hysteresis term due to our study using a 3-arm bandit versus a 2-arm bandit they used.

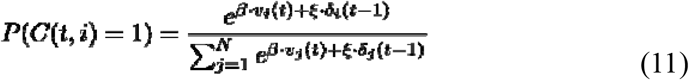

#### Exploration Bonus RL

The exploration bonus RL model was employed by Daw and Colleagues [27] in their 2006 investigation into human decision making, specifically to examine whether human exploration is directed towards the most uncertain actions. In their study, the model included a Kalman filter-based learning rule; however, we used the basic delta learning rule (Eq. 5) for consistency with other models in the comparison, as well as the fact that decision processes are the main focus here. The exploration bonus model’s decision rule is a modified SoftMax, Eq (12); adding the product of an “uncertainty bonus”, *E*_*j*_(*t*), and ϕ (∈ [0,1]) to the values in the exponential terms. *E*_*i*_(*t*) *and E*_*j*_(*t*) are counts of the number of trials since option i (or j) was last selected. Increases in these counts increase the probability of that option being chosen.

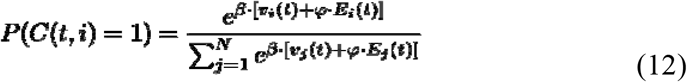

#### Model Comparison

Model best-fit (participant optimized) parameters were estimated for each participant for all models via MATLAB’s [28] fmincon function. All free parameters, see Table 2, were bound by definition except the inverse temperatures (βs). All βs span positive real numbers, however, in practice, no significant change in model output was seen once β ¿ 10. Therefore, for all simulations (including Model Generated Data below), β (∈ [0,10]). Multiple iterations were run for each participant’s choice data; during each iteration, model parameters were seeded with randomly generated initial values to avoid local minima. All value-comparison models’ optimization with fmincon was computed with switch or stay decisions on each trial and not the specific choice on a given trial in order for the foraging-RL model not to have an advantage in best-fit optimization.

**Table 2.** Summary and comparison of value-comparison RL models’ parameters and equations.

| Model | Learning Rule | Policy | Free Parameters |
| --- | --- | --- | --- |
| Dual Learning Rate | dual delta | basic SoftMax | $\alpha_{win}, \alpha_{loss}, \beta$ |
| Choice Kernel | value delta + ck delta | SoftMax w/ ck extended exponential | $\alpha, \beta, \alpha_{ck}, \beta_{ck}$ |
| Noisy Learning | delta w noise component | SoftMax w/ choice hysteresis | $\alpha, \beta, \zeta, \xi$ |
| Exploration Bonus | delta | SoftMax w/ eb value add | $\alpha, \beta, \varphi$ |

Each participant’s negative log-likelihood, computed in parallel with parameter optimization, was used to determine Akaike’s Information Criteria (AIC, [6]) values and Bayesian Information Criterion (BIC, [29]) values for model comparison. AIC and BIC weights were calculated according to Wagenmakers and Farrell [7].

#### Model Generated Data and Model Recovery

Outcome-sensitivity model generated results were the result of 10,0000 sets of simulations (artificial agent decisions) using randomly generated parameters drawn from uniform distributions. Each simulation set comprised fixed parameter values and 50 sessions (simulated bandits) of 300-trials. The random walks for the simulations used the same random walk parameters employed for the human participant task and each simulation set used one random walk for all 50 sessions. Lose-switch and win-switch rates were calculated for each session by dividing the total number of switches that occurred after each outcome by the total number of trials for each outcome. Lose-switch and Win-switch rates were then averaged across the 50 sessions.

Best-fit (participant-optimized) parameter model simulated data was generated in a similar manner, except the parameters for each set of 1000 simulations used the optimized parameters of a human participant. Model recovery used these best-fit generated data according to the procedure outlined in Wagenmakers et al. [9].

#### Behavioral (Self-Reported) Measures

4 participants out of 1000 were excluded from this analysis due to inconsistencies in the self-reported symptom measures. Each of the remaining participants’ model parameters were extracted and z-scored prior to further analysis.

A Poisson generalized linear model (GLM) was fit to a variety of outcome measures separately for each model’s parameter set. Specifically, for generalized anxiety symptoms and behavioral motivation, we first made a “best-parameter model” by fitting a univariate GLM to the one parameter that produced the highest *R*^2^ (e.g. inverse temperature (β) in the CK RL model for behavioral motivation) out of all parameters for both CK RL and foraging-RL model. We then constructed a “total-parameter model” by including all parameters from the CK RL model (α, β, α_*ck*_, β_*ck*_) or from the foraging-RL model (α, β, ρ) in a multivariate poisson GLM. We compared the *R*^2^ of the best-parameter model to that of the total-parameter model to see how much extra variance the full set of parameters explained.

We next extended the total-parameter model framework to eleven additional clinical and cognitive symptom scores, where we computed the Akaike Information Criterion:

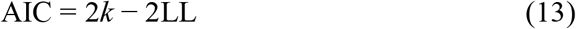

where *k* is the number of estimated regression coefficients (including the intercept) and *LL* is the Poisson log-likelihood evaluated at the fitted parameters. This permitted us to measure the AIC differences for each model relative to the best fit:

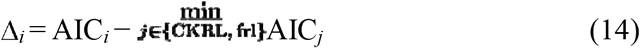

Finally, using these differences in AIC, we were next able to convert these values into the Akaike weights, which serves to estimate the relative likelihood of each model. This let us compare weights between the foraging-RL and CK RL models:

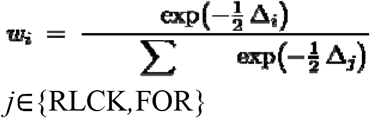

## Acknowledgments

This work was supported by NIMH under award #R21MH127607, NIDA under award #K23DA050909, and the University of Minnesota’s MnDRIVE (Minnesota’s Discovery, Research and Innovation Economy) initiative.

Yan, Xinyuan, R. Becket Ebitz, Nicola Grissom, David P. Darrow, and Alexander B. Herman. 2025. “Distinct Computational Mechanisms of Uncertainty Processing Explain Opposing Exploratory Behaviors in Anxiety and Apathy.” *Biological Psychiatry: Cognitive Neuroscience and Neuroimaging*, January. https://doi.org/10.1016/j.bpsc.2025.01.005.

## Footnotes

Note that if α_ck_ = 1 in the choice kernel learning rule of Eq (6), the choice kernel of the currently selected option will immediately go to one, while the choice kernel’s of the unselected options are immediately set to zero. Therefore, when α^ck=1^ choice kernel SoftMax and the choice hysteresis SoftMax rules are functionally equivalent (modulo β_ck=1_ and ξ

## Notes

### Competing Interest Statement

The authors have declared no competing interest.

### Summary of Updates

Minor changes to wording, including grammatical fixes, and changes to title. Updated the figures.

